# Animals in the Zika virus life cycle: what to expect from megadiverse Latin American countries

**DOI:** 10.1101/062034

**Authors:** Marina Galvão Bueno, Nádia Martinez, Lívia Abdala, Claudia Nunes Duarte dos Santos, Marcia Chame

## Abstract

Zika virus (ZIKV) was first isolated in 1947 in primates in Uganda, West Africa. The virus remained confined to the equatorial regions of Africa and Asia, cycling between infecting monkeys, arboreal mosquitoes, and occasional humans. The ZIKV Asiatic strain was probably introduced into Brazil in 2013. In the current critical human epidemic in the Americas, ZIKV is transmitted primarily by *Aedes aegypti* mosquitoes, especially where the human population density is combined with poor sanitation. Presently, ZIKV is in contact with the rich biodiversity in all Brazilian biomes, bordering on other Latin American countries. Infections in Brazilian primates have been reported recently, but the overall impact of this virus on wildlife in the Americas is still unknown. The current epidemic in the Americas requires knowledge on the role of mammals, especially non-human primates, in ZIKV transmission to humans. The article discusses the available data on ZIKV in host animals, besides issues of biodiversity, rapid environmental change, and impact on human health in megadiverse Latin American countries. The authors reviewed scientific articles and recent news stories on ZIKV in animals, showing that 47 animal species from three orders (mammals, reptiles, and birds) have been investigated for the potential to establish a sylvatic cycle. The review aims to contribute to epidemiological studies and the knowledge on the natural history of ZIKV. The article concludes with questions that require urgent attention in epidemiological studies involving wildlife in order to understand their role as ZIKV hosts and to effectively control the epidemic.

## INTRODUCTION

### A brief history of the Zika virus

Zika virus (ZIKV) is an emerging flavivirus from the same family as the West Nile (WNV), Japanese encephalitis (JEV), dengue (DENV), and yellow fever viruses (YFV) [1, 2]. ZIKV is an RNA virus, mostly transmitted to humans by bites from infected *Aedes* spp. mosquitoes, especially *Aedes aegypti*, which also transmits dengue and Chikungunya virus (CHIKV) in urban settings [3]. Other *Aedes* species have been implicated in ZIKV transmission, mainly in sylvatic cycles, including *A. africanus*, *A. albopictus, A. apicoargenteus*, and *A. furcifer* [4], 5, 6, 7].

ZIKV was first identified in 1947 in primates during a yellow fever virus study in Uganda [4]. The first reports of infected humans appeared five years later in Uganda and Tanzania [8], but the infection remained limited to equatorial regions of Africa and Asia, cycling between infective monkeys, arboreal mosquitoes, and occasional humans [9, 10]. Mosquitoes captured annually since 1965 in Senegal have shown that ZIKV amplifies cyclically every four years [11]. ZIKV outbreaks in humans occurred in 2007 on the island of Yap, Micronesia, and in Gabon [12, 5] and in 2013 in French Polynesia [13].

In the Americas, ZIKV is probably transmitted mainly by *Aedes aegypti*, a highly competent and anthropophilic vector species [14]. This mosquito, autochthonous to North Africa, spread to the Americas and Europe by the slave trade and adapted to the urban and domestic environment and, enabled the transmission of different arboviruses like DENV, YFV and CHIKV to humans, especially in areas with high population density and poor sanitation [15].

Species of the mosquito genera *Sabethes* and *Haemagogus* spp. have also been implicated in yellow fever transmission in the New World, and *Aedes albopictus*, which also occurs in the Americas, has been incriminated to transmitt ZIKV in Gabon [16]. However, the role of these vectors in maintaining ZIKV transmission in the Americas is not known.

Recent phylogenetic and molecular studies suggests a single introduction of the ZIKV Asiatic strain into the Americas (Brazil) between May and December 2013 [17] and in February 2014 in Chile [18]. In early 2015, several patients in Northeast Brazil presented dengue-like symptoms, and molecular diagnosis revealed autochthonous ZIKV infection [19].

An undetermined percentage of individuals with ZIKV infection fail to present clinical signs, but symptomatic individuals present mild fever, rash, headache, arthralgia, myalgia, asthenia, and non-purulent conjunctivitis three to twelve days after the mosquito vector bite [10, 13]. ZIKV infection poses a public health threat in Brazil, causing fetal microcephaly and other congenital malformations, besides other neurological disorders such as Guillain-Barré syndrome in adults [3].

ZIKV has invaded the huge biodiversity of all the Brazilian biomes, bordering on other Latin American countries. Althouse et al. (2016) ^[16]^ modeled the Zika virus transmission dynamics, estimating the numbers of primates and mosquitos needed to maintain a wild ZIKV cycle. Six thousand primates and 10,000 mosquitoes are enough to support a ZIKV transmission cycle. Based on the number of Brazilian primate species, the proximity of these and other small mammal species to urban and rural areas, and the wide distribution of *A. aegypti*, *A. albopictus*, and other mosquito genera like *Haemagogus* throughout the country, ZIKV spillover to wild primates is a potentially real scenario [20]. A wildlife cycle would launch new transmissions dynamics with unknown impacts on other animal species, including humans.

This review aims to describe the available data on ZIKV infection in host animals and its relationship to biodiversity, rapid environmental changes, and the impact on human health in megadiverse Latin American countries.

## METHODS

Recent advances in scientific research have emerged since ZIKV has become pandemic. We searched for scientific articles and news stories on research involving the ZIKV in animals using PubMed citation and index (http://www.ncbi.nlm.nih.gov/pubmed), the Fiocruz Library database (http://www.fiocruz.br/bibmang/cgi/cgilua.exe/sys/start.htm?tpl=home), Scopus database, (https://www.scopus.com) and websites for news stories in the mainstream lay press.

## RESULTS AND DISCUSSION

### Animals as ZIKV hosts

Few studies have focused on the role of animals as hosts for ZIKV. Some authors claim that there is no solid evidence of wild mammals such as non-human primates (NHP) as reservoirs for ZIKV. Meanwhile, studies have reported ZIKV antibodies in livestock like goats and sheep, rodents [21], and bats, lions, and ungulates like Artiodactyla, Perissodactyla, and Proboscidea [22]. According to a serological study in Senegal [23], monkeys from genera *Erythrocebuspatas* and *Clorocebussabeus* may act as ZIKV hosts in nature. In 1971, ZIKV antibodies were detected in primates from the Cercopithecidae family in Nigeria [24]. Several studies suggest that DENV, CHIKV and ZIKV adapted from an ancestral enzootic transmission cycle involving non-human primates and a broad spectrum of species from genus *Aedes* (*Stegomyia, aegypti*) as vectors in an urban-periurban cycle [25].

ZIKV infection has also been identified in naturally and experimentaly suscebtible other animal species (Table 1 and Fig. 1). Sera from 172 domestic animals and 157 wild rodents were tested for ZIKV in Pakistan, using the complement fixation test, showing that sheep, goats, some rodent species (*Tatera indica, Meriones hurrianae, Bandicota bengalensis)*, and one human living in the same area tested positive for ZIKV antibodies [21]. The authors suggested the need for a better understanding of this pathogen’s natural history.

**Fig 1.**
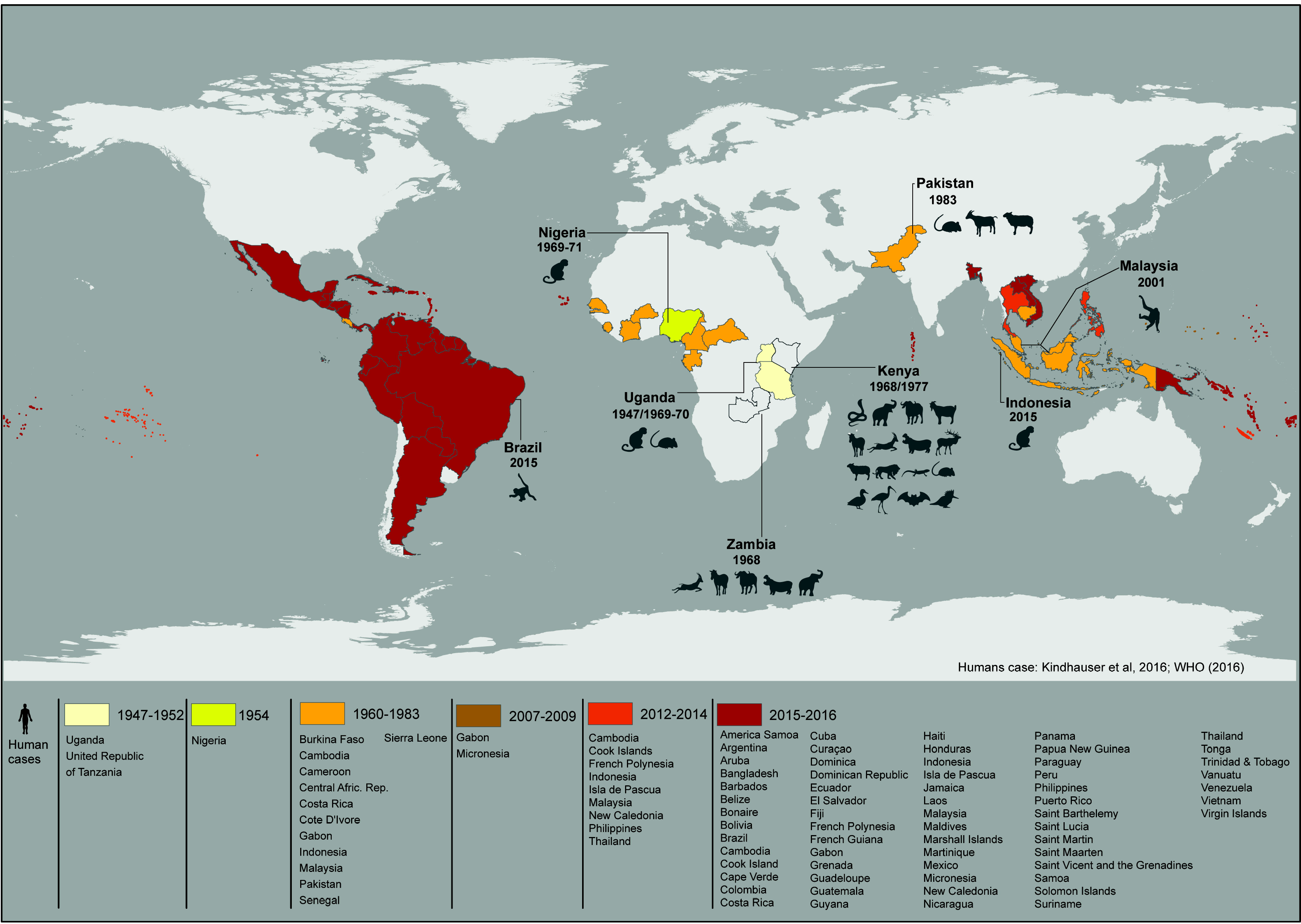
Historical time-line of Zika virus spread in humans and animals in the world. Colored countries have reported autochthonous vector-borne human cases, and those listed and with the years highlighted have reported diagnosed cases of ZIKV in naturally infected animals. The list of animal species is described in Table 1.

A study in Kenya in 1977 focused on the potential role of livestock (goats, sheep, and cattle) and wild vertebrates (2,424 small mammals, 1,202 birds, 18 reptiles) in maintaining arbovirus transmission. Hemagglutination inhibition assays showed that domestic animals (0.4%), wild birds (0.4%), small wild mammals (5.9%), and reptiles (27.7%) tested positive for ZIKV (Table 1) [26].

So, it is noteworthy that serologic studies should be interpreted carefully in view of possible cross-reactions with other antigenic flavivirurus, despite studies suggest that plaque reduction neutralization test (PRNT) do not cross and is the most specific serological test for the proper serological identification of flaviviruses [27, 28]

Unlike humans, wild mammals with ZIKV infection display few clinical signs. In a sentinel study in Uganda in 1947, primates showed only mild pyrexia. All monkeys inoculated by different routes developed neutralizing antibodies by day 14 after inoculation [4]. In the same study, Swiss mice became ill and one animal died following intracerebral inoculation [8]. Such inoculation is not a natural transmission route, and authors point out that some species of wild and laboratory rodents are resistant to some flavivirus infections, due to genetic resistance [29].

Most primates identified as ZIKV-positive in the wild or in sentinel studies are from Old World species. Phylogenetic analysis shows that humans are more closely related to Old World primate species, especially chimpanzees and orangutans [30]. Diseases that can be transmitted between closely related species often increase the relative risk. Non-human primates thus deserve special attention because of their close relatedness to humans and potential disease exchange [31].

Favoretto et. al. (2016) ^[20]^, using RT-PCR, showed that 29% of the New World primates *Callithrix jacchus* (common marmoset) *Sapajus libidinosus* (black-striped capuchin) in Ceará State in Northeast Brazil were infected with ZIKV. They also showed that the ZIKV genome sequence from monkeys was 100% similar to the ZIKV circulating in humans in South America, suggesting that primates sharing the habitat with humans could act as ZIKV reservoirs, as in the yellow fever sylvatic cycle in Brazil.

Besides the use of primates as sentinels in ZIKV studies, some experimental work has been performed with other mammals. Cotton-rats (*Sigmodon hispidus hispidus*), guinea pigs (*Cavia* sp.), and rabbits showed no clinical signs of infection after intracerebral inoculation [8]. An experiment in 1955 aimed to determine the susceptibility of cave bats (*Myotus lucifugus*) to ZIKV and showed that these bats are susceptible to ZIKV by intraperitoneal, intradermal, intracerebral, and intrarectal exposure, but not by intranasal exposure [32].

Barr et al. (2016) ^[33]^ infected cell cultures from different animal species with ZIKV and showed that 17 were susceptible to the virus, developing a cytopathic effect seven days postinfection. Some of the cell cultures were from domestic animals (dog, cats, chickens, horses, pigs, and cattle) and others from Old World wild animals (*Macaca mulatta*), while eight were from wild species found in the Americas: free-tailed bat (*Tabarida brasiliensis*), cottontail rabbit (*Sylvilagus floridanus*), gray fox (*Urocyon cinerorgeneus*), mule deer (*Odocoileus hemionus*), raccoon (*Procyon lotor*), Virginia opossum (*Didelphis virginiana*), nine-banded armadillo (*Dasypus novemcinctus*), Eastern woodchuck (*Marmota monax*), and American mink (*Neovison vison*). Most of these animals are peridomestic and sympatric to mosquito vectors. The authors also argued that with sufficiently high viremia, these animals could serve as reservoirs or hosts. However, they also indicated that the virus strain used in the experiment lacks some characteristics of the ZIKV currently circulating in the field, and that the virus in the laboratory does not mirror natural infection.

Public policy and elimination efforts in the Americas are currently based mainly on vector control and personal protection measures, so the high number of wild species with the potential to establish a sylvatic cycle makes elimination extremely difficult if not impossible [16]. We thus need studies on ZIKV in wild and domestic animals in the Americas, both to understand their potential role as hosts in the natural cycle and to target surveillance for enzootic ZIKV transmission.

### Biodiversity, animal hosts, and diseases

Human health relates closely to environmental health, defined here as the relationship between the health of domestic animals, wildlife, and the environment. Most etiological agents (60.3%) circulate between animals and humans (zoonotic diseases), and 71.8% of emerging diseases are caused by pathogens originating in wildlife [34]. A recent study associated 2,107 etiological agents with diseases in humans and animals [35].

Emerging diseases often occur in areas most heavily affected by natural events and human interventions, which further exacerbates social inequalities, health care costs, which influence the quality of life [36]. Vector-borne and parasitic diseases, with the disease burden driven by changes in biodiversity, have been shown to amplify the poverty cycle in many areas [37].

Recent efforts by the Convention on Biological Diversity and the World Health Organization have addressed scientific and political discussions on the relationship between human health and biodiversity. Such relationships include global concern over the importance of emerging zoonotic diseases originating in wildlife. Environmental changes, including loss of biodiversity, can favor emerging diseases originating from wildlife and act as the source of selective forces in new genetic variations leading to spillover and infecting humans [38]. This justifies actions to improve knowledge on biodiversity and pathogens and to monitor them to anticipate problems with installed competence. This approach has been strengthened by international and government programs that invest considerable resources in tracing pathogens worldwide. Monitoring diseases in animals poses a huge challenge for large, developing, and megadiverse countries like Brazil.

In this scenario, beyond seeking effective responses to health crises, we should implement measures that anticipate problems so that we can mitigate emerging diseases wherever possible and respond quickly when prevention and/or mitigation prove impossible or unfeasible.

The current ZIKV epidemic in Brazil requires understanding on the role of mammals, especially primates, in viral transmission to humans, especially when this interface occurs in fragmented forest areas as described by Favoretto et al. (2016) [20]. Such areas are usually bordered or surrounded by farmland and human settlements and by dense urban and unstructured areas that can increase contact between humans, wildlife, and domestic animals and occasionally promote disease spillover [39, 40]. Wild animals, especially primates, can thus be considered sentinels for pathogens of human health concern [41, 40]. ZIKV is an example of spillover, since this virus adapted from an ancestral transmission cycle involving non-human primates to an urban-periurban cycle, with humans as the main host.

Brazil is a megadiverse country with 357 million hectares of tropical forest and other highly biodiverse biomes. It is by far the world’s richest country in terms of biomes. Not surprisingly, Brazil has more primate species than any other country. Its 53 species account for 27% of the world’s primates. Forty of the 56 New World primate species are vulnerable, endangered, or critically endangered according to the IUCN red list of threatened species. The Atlantic Forest region is one of the highest priority areas for conservation in Brazil, since it is located in the most developed and most devastated part of the country [42], where 70% of the human population live between fragments of the natural forest [43].

Some non-human primate species occupy urban forests due to habitat fragmentation and have close contact with humans and domestic animals. Examples include primates from the Callitrichinae (*Callithrix, Leontopithecus*, and *Saguinus*), Cebinae (*Cebus*), and Atelidae families (*Alouatta* and *Brachyteles*) [44]. Favoretto et al. (2016) [20] were the first to report ZIKV in nonhuman primates in Northeast Brazil, highlighting that these New World primates can act as potential ZIKV reservoirs in the Americas. Many questions remain unanswered. Does ZIKV impact the health of non-human primates? Are NHPs living in urban fragments of forest more prone to ZIKV infection than those in preserved areas? Can naturally infected neotropical primates transmit ZIKV to mosquito vectors and thus help keep the virus circulating in the Americas?

Barr et al. (2016) [33] demonstrated the feasibility of infection in cell cultures from other mammalian species like carnivores, armadillos, rodents, and bats, thus raising the possibility of a transmission network shaped by biological and ecological factors. These include vector and host density and behavior, virulence, viral load, immunity, genetic variation, climate change, competition between biological communities, and anthropogenic forces like urbanization, sanitation, limited access to health services, poverty, and mistreatment of animals [29].

Considering the current epidemiological scenario with simultaneous circulation of the arboviruses ZIKV, DENV, and CHIKV and the fact that Brazil has a large non-human primate population, there is an urgent need to answer these questions to evaluate the impact of diseases like Zika on the non-human primate population in Brazil and elsewhere in the Americas. Yellow fever virus, another flavivirus that circulates in a sylvatic cycle in the Americas, has a great impact on primate populations, especially those of genus *Alouatta* [45] that exhibit disease signs after infection and act as sentinel primates for viral circulation and for implementation of control measures like human vaccination campaigns.

The pandemic ZIKV strain differs significantly from the African strain mainly in two regions of the genome. These acquired genetic markers increase its fitness for replication in the human host [3]. Whether these mutations also alter the infectivity in non-human primates remains to be determined. The role of wild primates and other mammals in ZIKV epidemiology thus requires urgent investigation.

Another relevant issue is the development of diagnostic tests for the detection of ZIKV infection in wild mammals, enabling unequivocal results without cross-reactivity with other flavivirus infections such as dengue and yellow fever.

A major threat to biodiversity is the introduction of invasive alien species (IAS) with potential impact on human health and infectious diseases. Some pathogenic parasites like mosquito-borne West Nile (also belonging to the genus flavivirus) virus can be categorized as IAS. Some authors consider modern pandemics like HIV and SARS as microbial-level invasion [38].

IAS can involve species, sub-species, or other taxa introduced by human action outside their natural (past or present) distribution and whose introduction, establishment, and spread threaten biological diversity and ecosystem integrity [38]. ZIKV is an example of IAS, due to it allocthonous origin and wide distribution in Brazil, and its origin in Africa, as with *Aedes aegypti* [46]. However, we do not know the impact of this virus on biodiversity, as with other IAS, or the ability of ZIKV to infect native vectors and mammalian hosts in the Americas or its potential to create new and different epidemiological cycles.

### Final Comments and Research Perspectives

Despite the growth of epidemiological knowledge in the last century, health interventions still mainly react to emergency events involving specific diseases in the human population, with some mitigation efforts [38]. The current ZIKV epidemic is no exception. We cannot expect to completely block the emergence of diseases, considering vector spread due to our limited capacity to reverse climate change, the globalization of goods and people, and our mode of production and consumption of natural resources. This situation is particularly paradoxical in megadiverse countries like Brazil.

The driving forces in the spread of diseases apply to the ZIKV epidemic, including anthropogenic activities, climatic change, intense human movement, loss of biodiversity, habitat destruction, land use change, introduction of invasive species, urban development, lack of knowledge on the role of animals in maintaining the sylvatic cycle, clinical manifestations, and wildlife trafficking [38]. The latter still occurs on a wide scale in Brazil. According to a national report, Brazil accounts for 5% to 15% of all smuggled animals in the world, with the removal of 12 million specimens from nature every year. Among animals trafficked in the New World, 95% are primate species from Brazil [47]. Wild animals are also extensively displaced inside Brazil due to domestic wildlife trafficking and human interventions like the construction of hydroelectric dams and highways. Such wildlife displacement has been implicated in increasing diseases and disseminating pathogens to new areas [48].

We need to understand the diversity of pathogens in nature and correlate them with biological communities, pathogenic and genetic characteristics, and anthropic impacts in areas where transmission and diseases occur. The complex evolutionary relationships between parasites, hosts, and vectors make this a challenging but strategically important task in face of the globalization, persistent poverty, and increasing environmental change. Awareness-raising is not enough to solve this problem. We need to expand the knowledge to diverse social actors and health and environmental services. The ZIKV epidemic illustrates the importance of monitoring and predicting the pathogens arising from wild animals and biodiversity.

Based on the above and the results of other studies, we pose several questions and hypotheses that emerge from this discussion and that require investigation:

1. What other wild animals besides primates could be infected by ZIKV in Americas? What is their role in maintaining and transmitting the virus to mosquito vectors? Which species can act as reservoirs?
2. Does the virus circulate at higher levels in wild animals inhabiting forest fragments adjacent to urban areas? What role do these animals play in maintaining the virus in areas close to humans?
3. Which wild hosts help keep the virus circulating in the Americas?
4. Do neotropical primates play a special role in the ZIKV epidemic?
5. Does the ZIKV impact wild animal populations and biodiversity? Does it cause disease and mortality in these animals?

Infectious diseases have important implications for animal and human health and biodiversity. Public health and biodiversity needs are misaligned and need to be rebalanced.

Rather than merely attacking and solving epidemic situations, as in the current ZIKV global health emergency, we need to predict and prevent future emerging diseases. Studies of wild hosts are troublesome and costly, especially when they require long-term monitoring. Funding also needs to be targeted for these studies. Future laboratory, field, and epidemiological research should focus on wildlife hosts to elucidate their role in ZIKV epidemiology in the Americas and enhance the epidemic’s control.

**Table 1.**
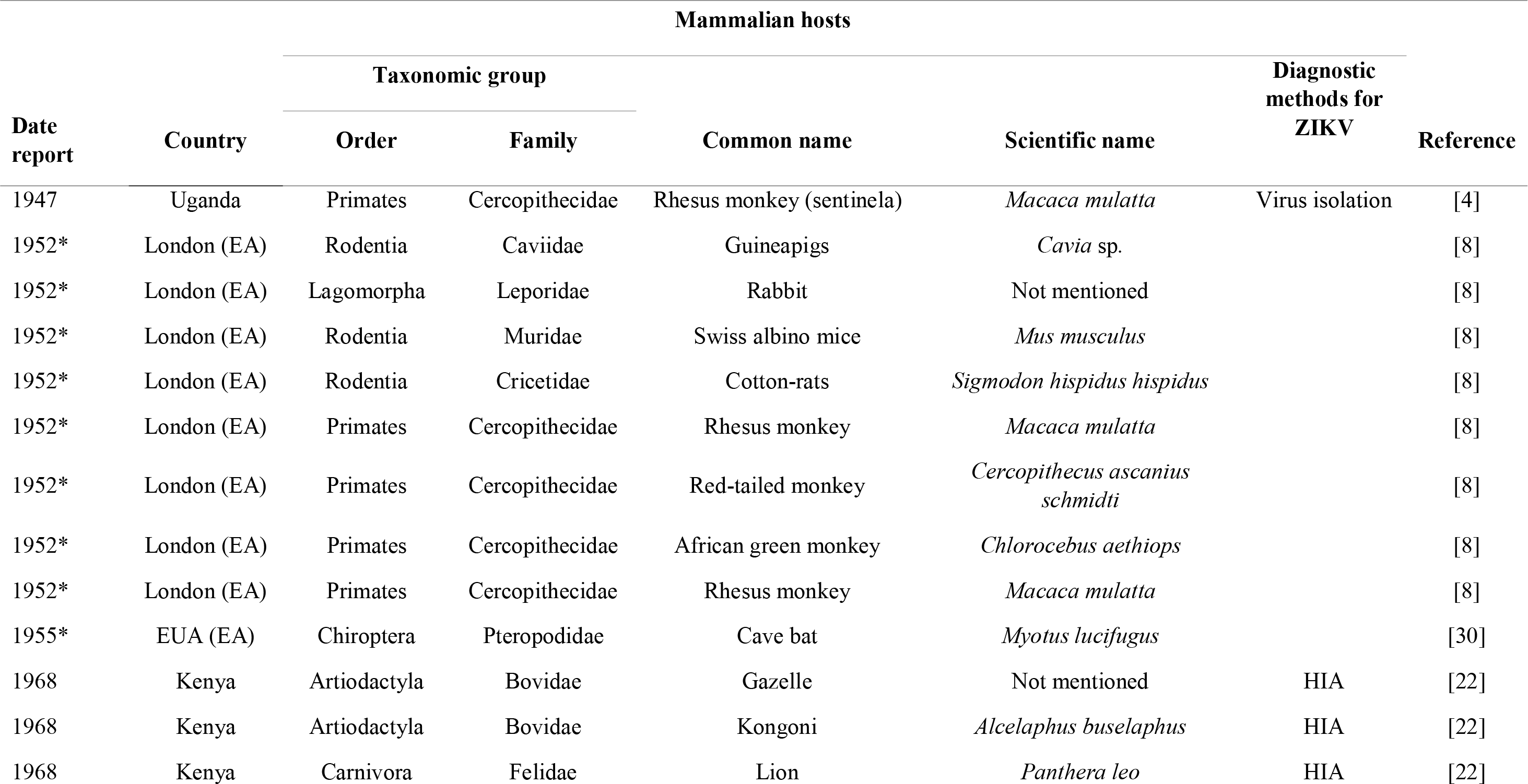
Chronological Zika virus natural and experimental assay (EA) infection in mammalian hosts in the world.

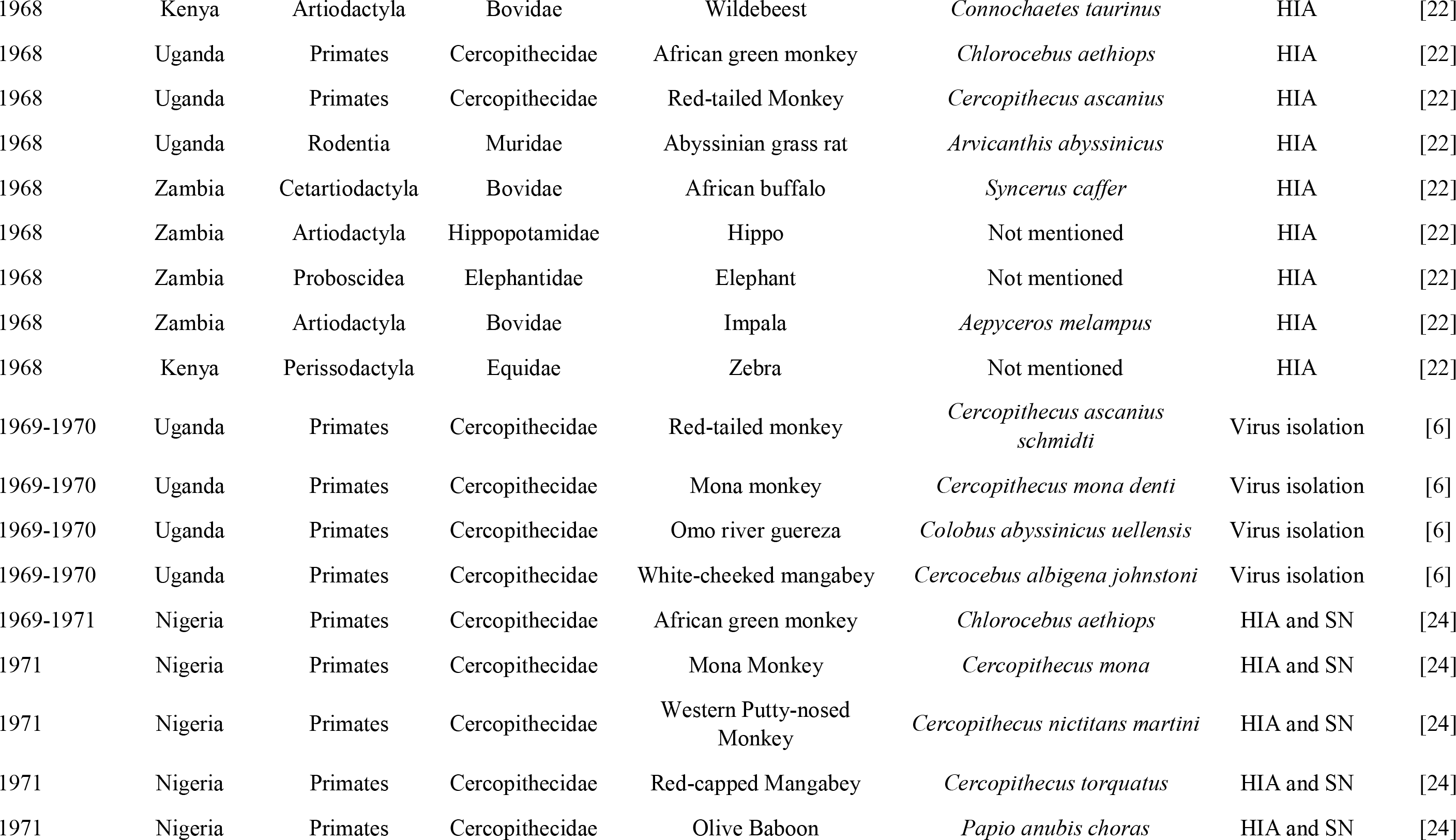

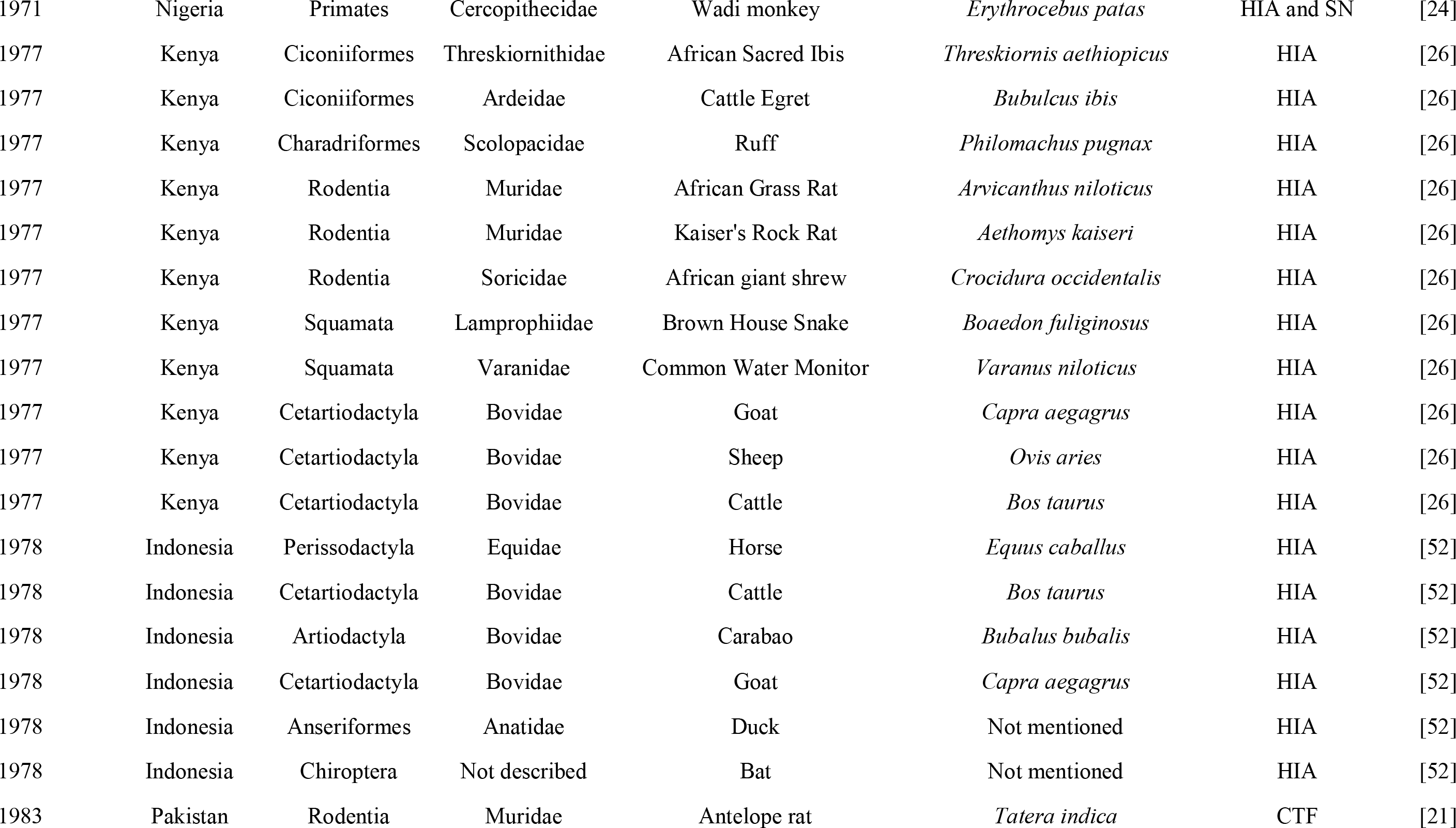

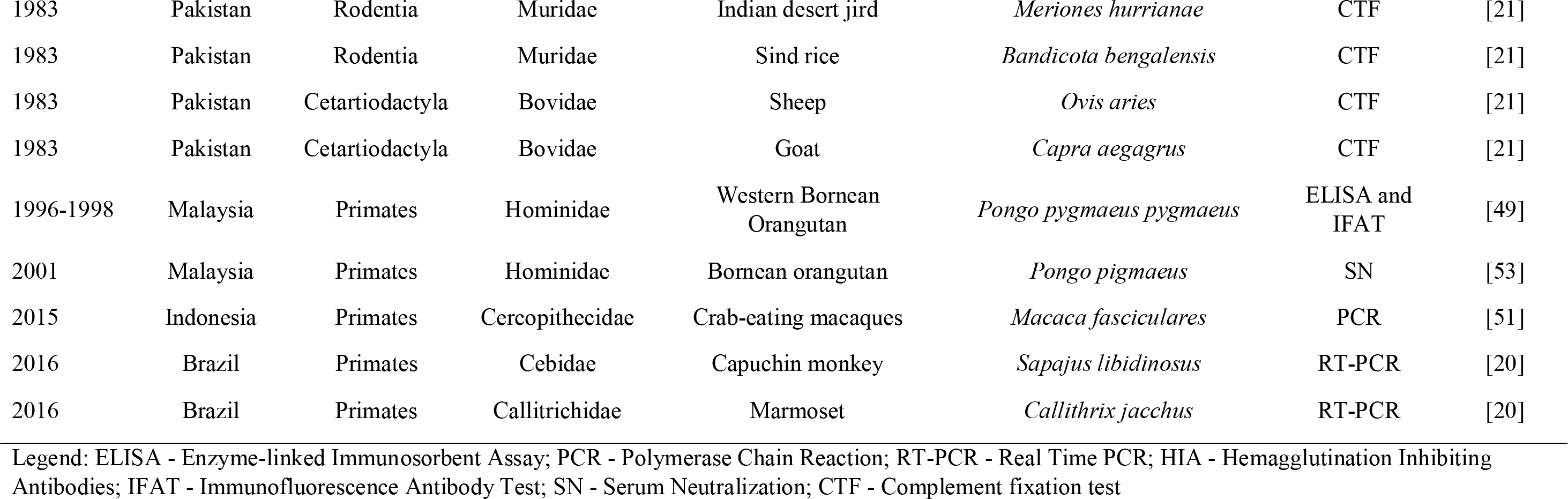

## Acknowledgments

We would like to thank the Fundafao Oswaldo Cruz (Fiocruz) for the support and we also thank Dr. Fernando Dias de Avila Pires for constructive comments.

